# Biotin based Northern Blotting (BiNoB): A Robust and Efficient Alternative

**DOI:** 10.1101/2024.07.08.602531

**Authors:** Deeksha Madhry, Riya Roy, Bhupendra Verma

## Abstract

Advancements in sequencing technology have led to the emergence of diverse types of regulatory RNAs. Differential transcript levels regulate cellular processes and influence disease severity. Identifying these variations through reliable methods is crucial for understanding their regulatory roles and disease mechanisms. Northern blotting, which used to be considered the gold standard for differential expression analysis, poses challenges due to various limitations associated with RNA quality, integrity, radioactivity, reagents, and the expenses associated with it. In this study, we employed a biotin-based Northern blotting approach, BiNoB, which offers advantages over traditional methods. In this study, we comprehensively targeted various RNA types, making this technique a versatile tool for RNA detection. Additionally, we conducted a comparison between 3’-end labeled probes, which were labeled in-house, and 5’-end labeled probes, which were commercially obtained. Remarkably, the results revealed significantly increased sensitivity with 3’-end labeled probes. Furthermore, we prepared an in-house buffer and compared its performance with the commercially available ULTRAhyb buffer, which exhibited comparable sensitivity, indicating that the in-house buffer is a cost-effective alternative. Intriguingly, our findings showed that as little as 1µg of total RNA proved to be adequate for the detection of small RNAs like tRNAs and their derived fragments.

## Introduction

Recent progress in deep sequencing and high-throughput screening has significantly improved our understanding of the RNAome, constantly transforming our insights into small noncoding RNAs (sncRNAs). These includes miRNA, siRNA, tRNA-derived RNA fragments (tsRNA), Piwi RNA, and other categories collectively known as sncRNA. Despite their lack of protein-coding ability, these sncRNAs play diverse regulatory roles, finely tuning host gene expression at epigenetic, transcriptional, and posttranscriptional levels. Their modulation is pivotal for various cellular pathways such as the cell cycle, apoptosis, transcription, translation, cell proliferation, and differentiation. The differential expression of these sncRNAs can serve as crucial biomarkers for identifying disease severity which has necessitated the precise detection and quantification of their expression levels using a standardized and reliable method.

The traditional northern blotting method is still considered as the gold standard as it has the advantage to visually inspect and provide approximate quantification of the expression levels. However, in the previous years, it was considered to have less sensitivity without radioisotope labelled probes, consequently making it unsuitable for large scale expression analysis. To address these limitations, diverse approaches such as microarray-based, PCR-based, and sequencing-based methods have emerged ^1–5^. While these techniques have significantly enhanced the quantification of small non-coding RNAs (sncRNAs), they have the disadvantage of not elucidating their size distribution, a crucial aspect for studying their biogenesis. Hence, northern blotting has regained significance, offering potential for advancements by enhancing sensitivity and detection capabilities regarding the size distribution and biogenesis of small RNAs.

Different studies have reported varied improvements in the northern blotting technique, primarily based on the method of probe labeling-radioactive and non-radioactive. Traditional northern blotting is based on the 5’end radio-labelling of DNA probes using γATP mediated kinase reaction. This approach enhances the detection sensitivity for less abundant transcripts, resulting in an improved signal-to-noise ratio. However, the use of radioisotopes has many shortcomings 1) it requires a dedicated facility for handling and disposal, 2) specialized training is needed to work in radioactive facility, and 3) the limited half-life of radioisotopes restricts their use for shorter duration only. In contrasts, non-radioactive probe labelling methods such as DIG (Digoxigenin), biotin, and fluorescence offers the convenience of easy handling and storage for longer durations, making them cost effectivealternatives^6–8^. Figure 1 shows the differences between the mechanisms of radioactive and non-radioactive probe labelling.

**Figure 1:**
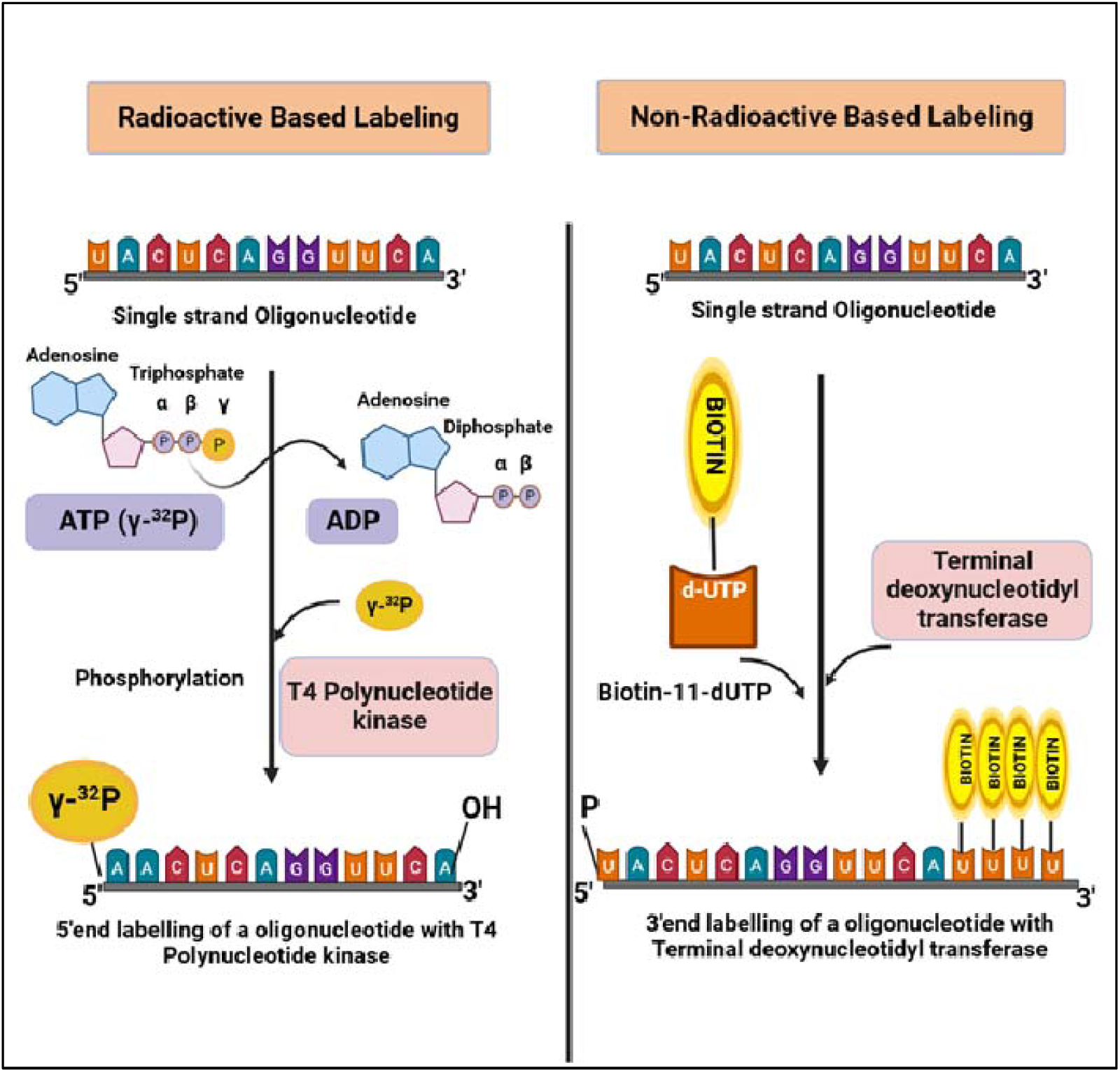
Comparison between mechanisms of radioactive and non-radioactive based labeling: Radioactive-based labelling utilizes the T4 polynucleotide kinase enzyme, which catalyzes the transfer of a phosphate group from ATP to the 5’ end of RNA oligos. A radioactive phosphate, often ^32^P, is incorporated into the 5’ end of probes during this process. The non-radioactive approach involves labelling probe with biotin-11-dUTP at the 3’ end of the RNA strand with the help of terminal deoxynucleotidyl transferase.

Secondly, Locked Nucleic Acid (LNA) based probes represent another improvement that provides better sensitivity than DNA probes. LNAs form a bridge between the 2’-O, and 4’-C methyl group of ribose sugar, stabilizing the ribose sugar conformation. This elevates the thermal stability of the LNA oligos, resulting in increased affinity binding between the RNA-probe duplex and ultimately enhancing the sensitivity of small RNA detection ^9–11^.

Thirdly, the RNA immobilization method has also been improved for the increased sensitivity. Previously, RNA was immobilized onto the solid support of nylon membrane by either baking at 80°C or by UV crosslinking. UV crosslinking of the membrane is selective, primarily facilitating crosslinking between the free amino groups of the membrane and Uracil nucleotides of the RNA. This selectivity limits crosslinking across the entire RNA length and introduces steric hindrance, reducing accessibility to immobilized RNA. These factors pose limitations, particularly for small RNAs, making their immobilization challenging and restricting accessibility due to size constraints. Consequently, this process tends to be less sensitive for small hybridization, impacting the overall detection efficiency. An effective cross-linking method for small RNA detection was devised by Pall et all. using water soluble EDC (1-ethyl-3-(3-dimethyl aminopropyl) carbodiimide) chemical crosslinking ^12^. EDC helps in crosslinking of free amine groups of membrane with the 5’ phosphate group of RNA, resulting in reduced steric hinderance and an increase in probe binding efficiency ^13,14^.

Despite numerous advancements in the Northern blotting technique, its regular use is hindered by challenges such as requirement for handling expertise, cost, and the need for specialized setups. In this study, we aimed to establish a ‘NO FAIL’ protocol that addresses these concerns. We have optimized **<u>Bi</u>**otin-based **<u>No</u>**rthern **<u>B</u>**lotting (BiNoB) protocol to enhance sensitivity across a range of RNA lengths in a cost-effective manner. This protocol eliminates the need for special setups, expensive reagents, or specialized training. Notably, we successfully applied this approach to detect small RNAs as short as 22 nucleotides, as well as large RNAs spanning kilobases in size.

## Materials and Methods

### RNA Extraction

Total cellular RNA was isolated from different samples using the TRIzol method. Briefly, 300 µl of chloroform was added to 500 µl of TRIzol extracts, followed by thorough mixing by inverting the tubes and incubate in at room temperature for 10 min. The tubes were then centrifuged at 13,000 rpm for 10 min at 4°C to collect the aqueous supernatant. To this supernatant, 600 µl of chilled absolute ethanol was added, thoroughly mixed, and then passed through RNA columns at 8000 rpm for 30 seconds. The column was subsequently washed with AW1 and AW2 buffers (Qiagen). Following 1min of empty spin, the RNA was finally eluted in nuclease-free water. RNA concentration was measured using Nanodrop 1000 and absorbance ratio of 260/280 and 260/230 nm were checked for RNA quality. All the chemical reagents used here are mentioned in supplementary 1.

### Sample preparation and Gel electrophoresis

RNA samples were mixed with 2x Urea-loading dye (90mM Tris Base, 90 mM Boric acid, 2 mM EDTA, 12% Ficoll, 7M Urea, 0.03 %Xylene cyanol and Bromophenol blue dye) in equal volume, followed by heat treatment at 95°C for 5 minutes.

A 12% Urea-PAGE gel was prepared for both high molecular weight (more than 2Kb) and low molecular weight (40 bp-22 bp) RNA. The gel composition used was 6 ml of 20% acrylamide:bis-acrylamide (19:1) solution, 4 ml of 45% urea buffer prepared in 1x TBE (Tris Borate EDTA) buffer, 50 µl of 10 % APS, and 10 µl TEMED. The gel was pre-run for 45 minutes at 150 V in 1x TBE buffer. Samples were then loaded and run the gel at 150 V till the bromophenol dye reached lower part of the gel. Subsequently, the gel was stained with EtBr (0.05 µg/ml) for 15 min followed by its visualization on the ChemiDoc system (Syngene G:Box system).

### Membrane transfer and crosslinking

Wet transfer was performed. Briefly, whatmann paper, sponges and the Hybond N+ Nylon membrane (Cytiva: RPN203B) were presoaked in the transfer buffer (0.5x TBE). Wet transfer was carried out for 1hr 30min at 100V in 0.5x TBE buffer in mini trans-blot cell Bio-Rad apparatus. Following transfer, the membrane was UV crosslinked in UV crosslinker (Analytik jena) at 1250 mJ/cm^2^. Baking was performed for 2 hours at 80°C after UV crosslinking.

### Probe labelling and dot blot analysis

DNA probes were biotinylated at the 3’terminal using Biotin 11-dUTP (Thermo: R0081) and terminal deoxynucleotidyl transferase TdT enzyme (Thermo scientific: EP0161) in the presence of CaCl2 containing 5X reaction buffer, according to the manufacturer’s protocol. Biotin labeling reaction has been provided in table 2. The reaction was incubated at 37°C for 15 min followed by heat inactivation at 70°C for 10 min. After completion of the reaction, the biotinylated probe mixture was passed through a G-25 spin column (Cytiva: 27532501) by centrifugation at 13,000 rpm for 1 minute at room temperature to remove unincorporated biotin moieties. Also, 5’end biotinylated probes were commercially purchased from Bioserve Biotechnologies (India) Pvt.Ltd to compare the sensitivity of both the probes. The sequences of probes used in the study has been mention in the table 3.

**Table 1:** A Comparison between Classical Northern Blotting and the BiNoB protocol.

| **Step** | **Classical method** | **BiNoB method** |
| --- | --- | --- |
| **Gel preparation** | • Formaldehyde/ polyacrylamide gel.<br>• Larger volume of gel required ~200ml. | • Polyacrylamide gel<br>• Less volume is required ~15-20ml. |
| **Electrophoresis** | • Horizontal gel electrophoresis<br>• 100V for ~2-3hr. | • Vertical gel electrophoresis<br>• 120-130V for ~1hr. |
| **Gel transfer** | • Capillary method.<br>• Overnight transfer for 12-16hrs. | • Wet transfer.<br>• 100V for 90min duration |
| **Probes used** | • DNA or RNA | • DNA and LNA probes |
| **Probe labelling** | • Radioactive labelling<br>• Limited access | • Non-radioactive labelling.<br>• Accessible to all. |
| **Crosslinking** | • UV crosslinking | • UV crosslinking followed by baking at 80°C. |
| **Signal detection** | • More than 8hrs of exposure is required on phosphoscreen and scanned using phosphoimager. | • Rapid detection<br>• Using ECL kit and ChemiDoc system |

**Table 2:** Biotin labelling reaction.

| **Components** | **Volume/ concentration** |
| --- | --- |
| **5X reaction buffer for TdT** | 10 µl |
| **DNA oligos** | 10 pmol |
| **Biotin-11-dUTP (1mM)** | 4 µl |
| **TdT** | 40 U |
| **Nuclease free water** | to 100 µl |

**Table 3:**
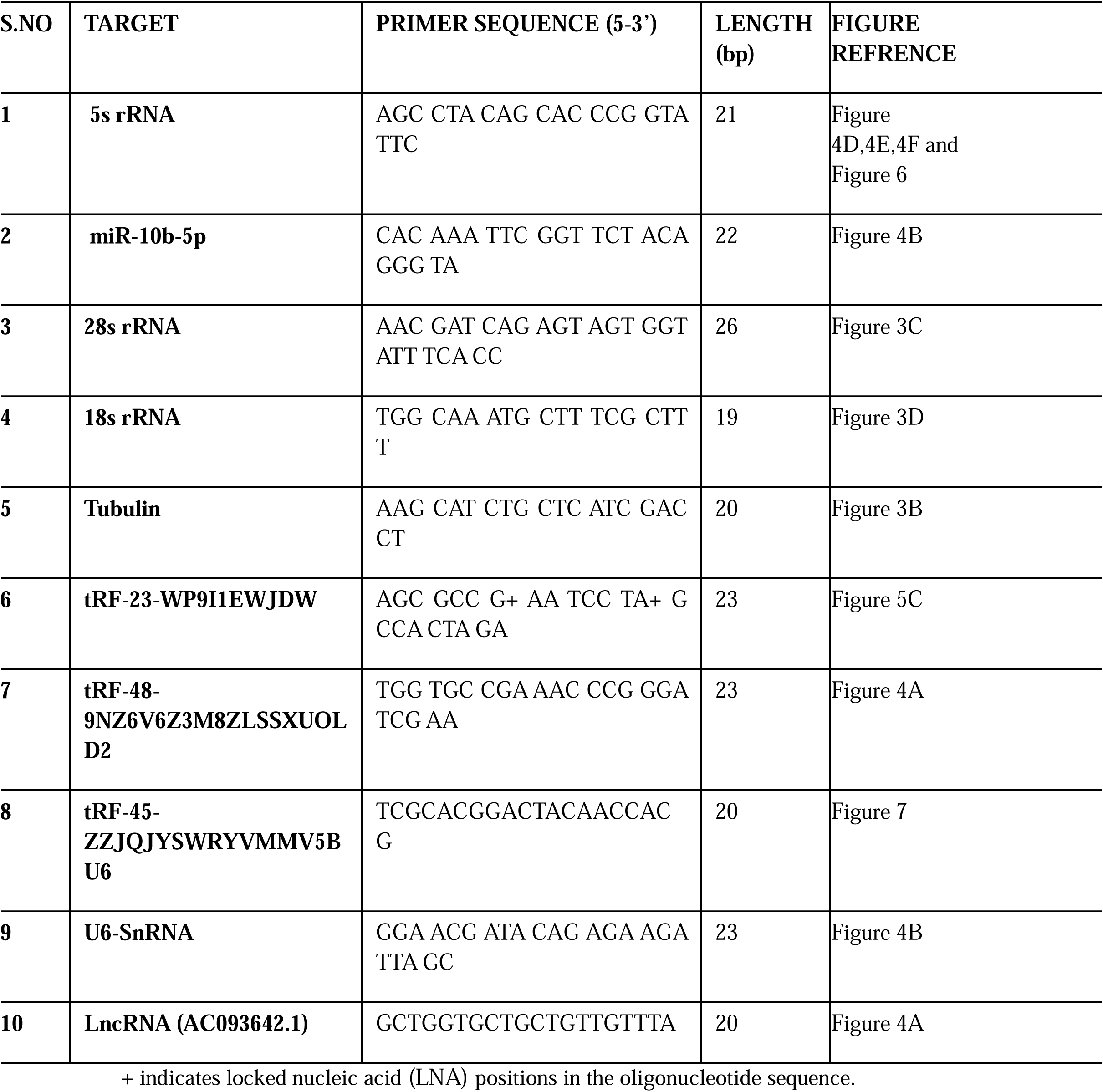
Sequences of probes utilized for the northern blot.

Biotinylation of probes was confirmed through dot blot analysis. Two microliter of each biotinylated probe was spotted onto the pre-soaked Hybond N+ Nylon membrane in 1x TBE buffer. The membrane was then UV-cross-linked at (1250 mJ/cm2) followed by incubation in blocking solution (0.1% skimmed milk in 1X TBS with 0.1% tween 20) for 30 minutes at RT. The membrane is finally incubated with streptavidin-HRP conjugate at RT for 20 minutes, followed by washing with low-stringency buffer (2X SSC + 0.1% SDS) and high-stringency buffer (0.1% SSC + 0.1% SDS) for 5 minutes each.

### Prehybridization and Hybridization

Membranes were pre-hybridized for 3 hours at 50°C in an in-house prepared buffer (10mg/ml of BSA, 0.5 M sodium phosphate buffer (pH 7.2), 1mM EDTA,7% SDS). Following this, overnight hybridization was set up for 12-16 hours by adding 25 µl of the biotin-labelled probes to the same buffer at 50°C. The commercially available ULTRAHyb^TM^ buffer (Ambion:AM8670) was also used for comparison with the in-house-prepared buffer.

### Membrane blocking and Streptavidin binding and washing

Following hybridization process, the membrane was first washed with a low stringency buffer (2X SSC+ 0.1% SDS), then by high stringency buffer (0.1% SSC+0.1% SDS) for five minutes each. Subsequently, the membrane was subjected to a blocking solution (0.1% skimmed milk in 1X TBS with 0.1% tween 20) for 1 hour at room temperature. Next, the membrane was incubated with HRP-Streptavidin conjugate at room temperature for 30 minutes, followed by a final washing of 30 minutes in washing solution (1X TBS + 0.1% SDS).

### Signal detection

After the final washing step, the blot was finally developed using ECL reagents (Thermo:32106) and the blot was finally visualized on ChemiDoc system (Syngene G: Box system).

### Membrane stripping

The membrane was stripped overnight at 60°C in a stripping buffer containing 1.5% SDS (sodium dodecyl sulfate) prepared in autoclaved distilled water.

## Results

### Biotin labelling of probes

To detect RNA using northern blotting, radioactive-based labeling was often used previously. However, due to its inconvenient and hazardous process, it has been replaced by many non-radioactive labels like DIG (digoxigenin) and biotin to increase the sensitivity and specificity of small RNAs for detection. Biotin has high-affinity binding to streptavidin, which is the strongest of any non-covalent bond. This affinity is significantly higher than that between DIG and an anti-DIG monoclonal antibody, contributing to the greater reproducibility of the biotin/streptavidin system.^15^ For our study, we therefore, have designed probes labeled with biotin to detect the bands of large and small RNAs. This method is very sensitive and capable of detecting even small sized RNA, about 22 nt in length. Biotin labeling can be performed at both ends of the oligonucleotide, either at the 5’ end or the 3’ end terminus^16^. In our studies, both labeling techniques have been utilized to compare the sensitivity and detection of target RNAs. These labeled probes are stable and can be stored at −20°C for long-term purposes ^16^.

### Dot blot confirmation

In order to check the biotin labeling of selected probes, dot blot hybridization was performed. In our study, we performed 3’ end biotin labelling of probes for the detection of various RNAs (5s rRNA, 28s rRNA, 18s rRNA, tRNA, tRF 48 (tRF-48-9NZ6V6Z3M8ZLSSXUOLD2), miR-10b-5p and tRF(tRF-23-WP9I1EWJDW). The labelled probes were spotted onto membrane and then UV crosslinked, blocking and washing steps were carried out, followed by exposure to an imaging system. A dark dot observed in the blot indicates proper biotin labelling (figure 2).The dot blot of the labelled probes confirms the suitability for further northern hybridization detection.

**Figure 2:**
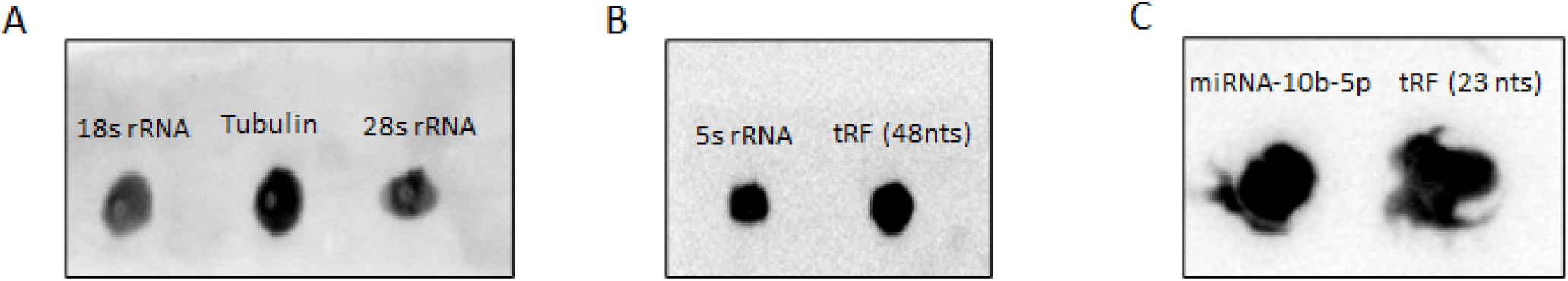
Dot blot Assay: Different probes were labelled with biotin-11-dUTP using the enzyme Terminal deoxynucleotidyl transferase. The biotin labelling of the probes was confirmed by dot blot analysis prior to membrane hybridization.

### RNA Quality and integrity check

RNA was isolated using TRIzol extracts from three different cell lines: Huh7 orginated from human hepatocellular carcinoma, HEK from human embryonic kidney cells and Vero kidney epithelial cells extracted from an African green monkey were marked as RNA1, RNA2 and RNA3, respectively. RNA quality and integrity are the primary factors for northern blot analysis. Our results for RNA1, RNA2, and RNA3 showed a ratio of absorbance at 260 and 280 nm in the range of 2.0-2.2, which ensures high purity and yields a higher concentration of RNAs. RNA integrity was also assessed by Urea-PAGE gel electrophoresis followed by EtBr staining. EtBr staining revealed clear bands of five significant RNA species, with 28 and 18S RNAs (ranging from 2.8Kb to 1.8Kb in size) observed exclusively in the wells, followed by 5.8s rRNA (160 nucleotides), 5s rRNA (125 nucleotides), and tRNA (80–100 nucleotides) (figure 3A).

**Figure 3:**
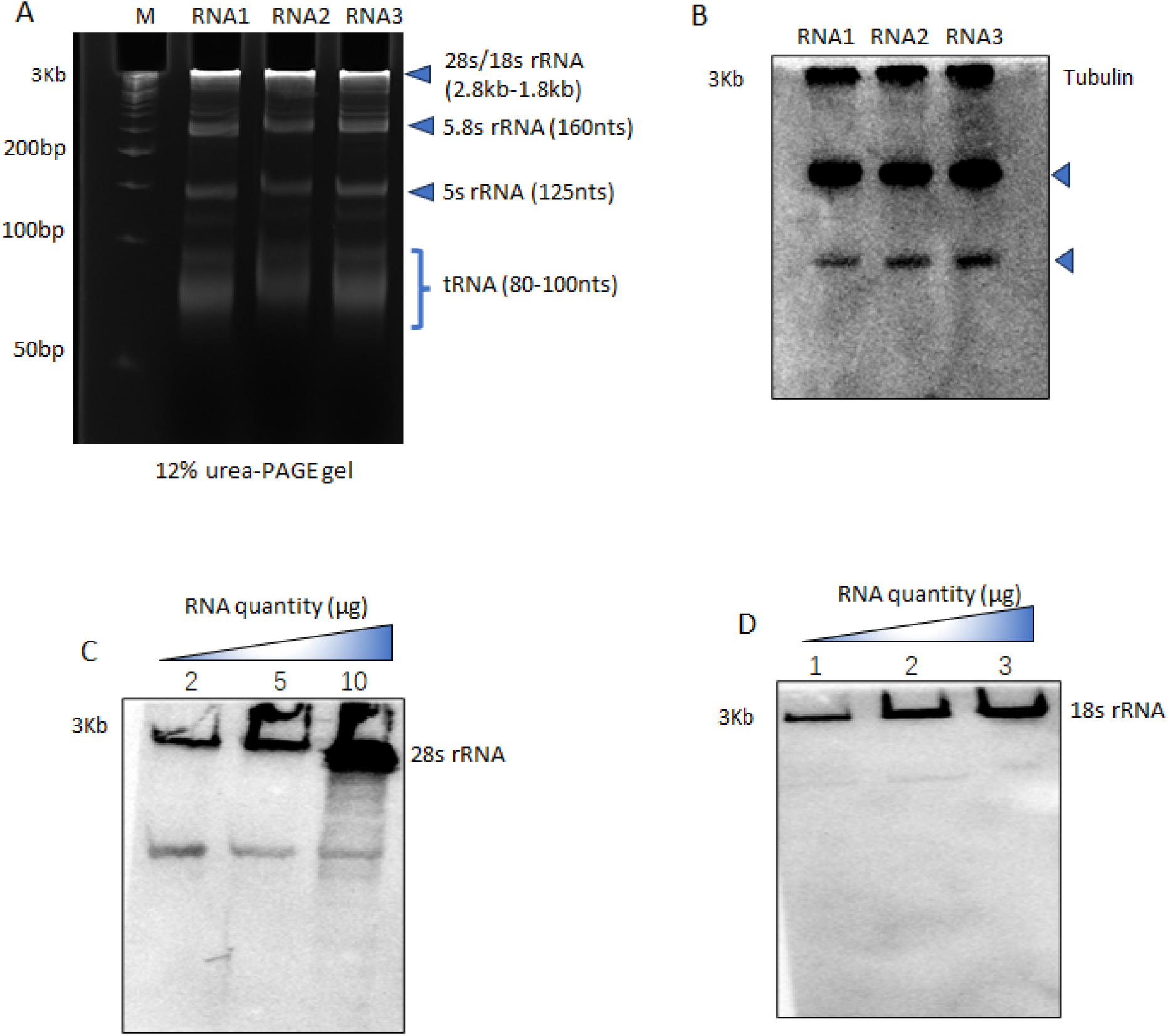
RNA integrity check and large sized RNA detection: Total RNA (RNA1, RNA2, and RNA3) was extracted from the Huh7, HEK and Vero cell lines, and 5 µg of total RNA was loaded into a 12% urea-PAGE denaturing gel. A) EtBr staining was performed to check the RNA quality and integrity. Then, the RNA samples were transferred onto a nylon membrane. Subsequently, the membrane was hybridized with biotin labeled probes targeting B) tubulin. (The lower bands marked with blue triangles, correspond to 5S rRNA and tRNA, for which the membrane was initially probed before probing for tubulin.) Various RNA quantities used to determine the sensitivity of detection. C) RNA quantity used for the detection of 28S rRNA were 2µg, 5µg, and 10µg, while for D) 18SrRNA, RNA quantity was further reduced to 1µg, 2µg, and 3µg. (M: 50bp ladder).

### BiNoB detection of Long-sized RNAs (28s rRNA,18s rRNA and tubulin)

To efficiently detect large sized RNAs with more than 1kb in size, specific probes were designed and biotinylated at the 3’-end using TdT (Terminal deoxynucleotidyl Transferase) enzyme. RNA was loaded onto a 12% urea denaturing PAGE, followed by EtBr staining to analyze the RNA quality and integrity. The gel was then transferred onto the Hybond nylon membrane though wet transfer, followed by membrane UV crosslinking and baking. The membrane was subsequently hybridized to detect large sized rRNAs and mRNAs in the kilobase pair range. The sequence specific probes against tubulin, 28S rRNA and 18S rRNA were used for hybridization. Tubulin was detected in different cell lines; Huh7, HEK cells and Vero marked as RNA1, RNA2 and RNA3, respectively, with 5ug of each RNA sample was used for the detection. The results showed clear bands of tubulin in all the cell lines (more than 1kb) around the wells in equal intensity (figure 3B).

Next, to determine the sensitivity of the technique for the detection of large sized RNAs, different quantities of RNA ranging from 1-10 µg were used. For the detection of 28S rRNA, RNA quantities used were 2 µg, 5 µg, and 10 µg (figure 3C). while for 18SrRNA, quantities were further reduced to 1 µg, 2 µg, and 3µg (figure 3D). The results demonstrated a clear and concentration-dependent increase in band intensity for both 28S and 18S rRNA. These results suggest that a quantity of 1 µg is sufficient for detecting large-sized RNAs, indicating the technique’s sensitivity and specificity.

### BiNoB detection of Mid-sized RNAs (Lnc RNA, 5s rRNA, U6snRNA and tRNA)

We next aimed to detect the RNAs ranging from 80-600bp in length. For this purpose, 5µg RNA samples from Huh7, HEK cells, and Vero cells, labeled as RNA1, RNA2, and RNA3 respectively, were prepared. Various types of RNA, including long non-coding (lnc) RNA AC093642.1 (∼600 nucleotides), U6 snRNA (∼100 base pairs), and tRNA-Phe GAA (80-100 base pairs), were targeted using 3’ biotinylated probes. Clear single bands corresponding to lncRNA and tRNA were observed, while two bands were detected for U6, possibly due to multiple U6 transcripts (figure 4A, 4B, 4C). Additionally, 5s rRNA, with 125 nucleotides in length, was detected across different RNA quantities (1 µg, 2 µg, and 3 µg), showing clear single bands exhibiting increasing intensity with increasing RNA quantity (figure 4D).

**Figure 4:**
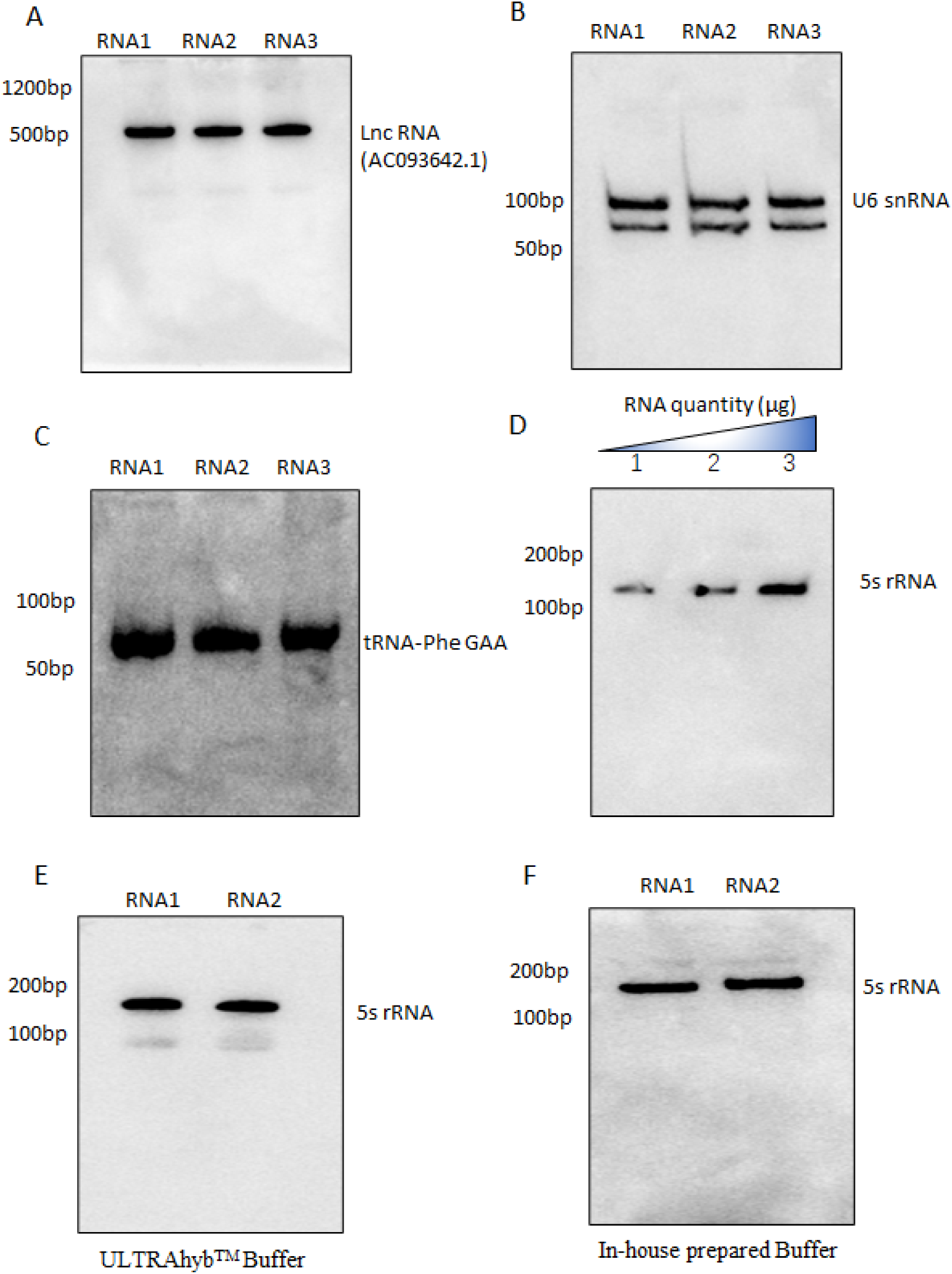
Mid-sized RNA detection: RNA samples (5µg) were loaded into 12% urea PAGE followed by transfer onto nylon membrane and hybridized with different biotinylated probes. A) lncRNA having ∼600 nucleotides, B) tRNA, and C) U6 snRNA, which are approximately 80-100 nucleotides. D) depicts the bands corresponding to 5s, which are about 125 nucleotides. The comparison between two different hybridization buffers E) ULTRAhyb^TM^ ultrasensitive hybridization buffer and F) an in-house prepared buffer was used for the detection of 5s rRNA.

A comparative analysis was also performed to observe the sensitivity of detection using ULTRAhyb^TM^ ultrasensitive hybridization buffer (Invitrogen) and in-house prepared hybridization buffer for the detection of 5s rRNAs (figure 4E). RNA extracts from Huh7 and Vero cells (RNA1 and RNA2) were used, with each sample containing 5 µg of RNA. The results showed a single clear band of 5s rRNA with comparable sensitivity. Our results suggest that the in-house prepared buffer demonstrates the same level of sensitivity compared to that of ULTRAhyb buffer.

### BiNoB detection of Small non-coding RNAs (miRNA and tRNA fragments)

For the detection of small RNAs, our objective was to detect miRNA and tRNA-derived fragments (tRFs), which typically range from approximately 18-50 nts in length. Five micrograms of each RNA sample were utilized for small RNA detection. The sequence specific probes were used for the detection of miRNA (miR-10b-5p) which is 22ntd in length, and a tRNA fragment derived from tRNA-Phe-GAA (MINTbase ID: tRF-48-9NZ6V6Z3M8ZLSSXUOLD2). In figure 5A, two clear specific bands of tRNA, approximately 80-100 ntd, and its derived fragment, 48 ntd in length, were observed. In the case of miRNA (figure 5B), one clear band at approximately 22 ntd and two diffused bands at around 100 and 1000 ntd were observed. The biogenesis of miRNAs involves two RNA cleavage stages. Initially, within the nucleus, the ribonuclease Drosha forms a complex with the protein DGCR8 to cleave miRNA transcripts (pri-miRNA), which are ∼1000 ntd long, into pre-miRNA precursors, ∼60-120 ntd in length. Subsequently, these pre-miRNAs are transported to the cytoplasm by Exportin-5, where they undergo further processing by the ribonuclease Dicer protein complex. This secondary cleavage results in the formation of miRNA duplexes ∼20-30 ntd in length ^17–19^. This sensitivity of the BiNoB technique enabled us to detect the three forms of miRNA: pri-miRNA (1000 nucleotides), pre-miRNA (60-120 nucleotides), and miRNA (20–25 nucleotides). These findings suggests that the BiNoB technique is suitable for studying miRNA biogenesis.

**Figure 5:**
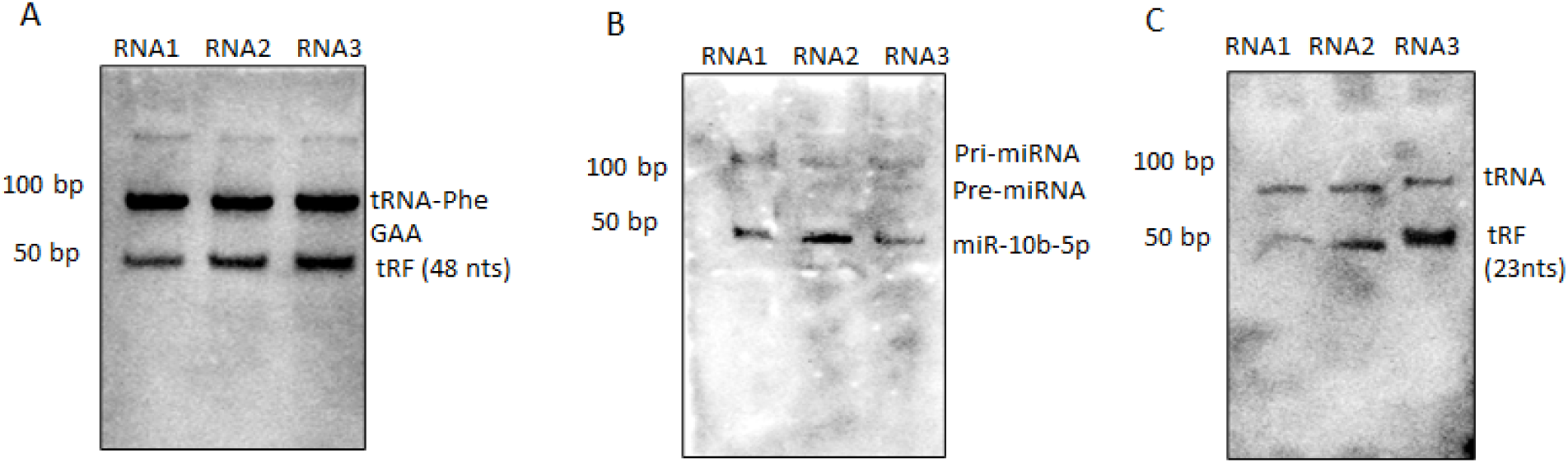
Detection of small sized RNA using BiNoB: Total RNA samples (5ug) were loaded onto a 12% urea denaturing gel and transferred into a nylon membrane, followed by hybridization with biotin labeled probes. A) The blot represents the presence of a tRF (9NZ6V6Z3M8ZLSSXUOLD2) with a length of 48 nucleotides, and its parent tRNA (Phe-GAA). B) miRNA detection demonstrated the successful detection of three forms of miRNA: pri-miRNA (1000 nucleotides), pre-miRNA (60-120 nucleotides), and miR-10b-5p (20–25 nucleotides). C) LNA modified probe was used to detect small length tRNA fragment of 23 nucleotides in the blot using BiNoB approach.

### BiNoB detection using LNA probes

We also utilized LNA (Locked Nucleic Acid) based probes to compare the sensitivity and specificity of DNA versus LNA probes. A sequence specific LNA probe was used for the detection of specific tRNA-fragment of 23ntd in length, derived from tRNA-GluTTC (MINTbase ID: tRF-23-WP9I1EWJDW). We distinctly observed two bands corresponding to tRNA (∼100ntd) and its fragment (23ntd) (figure 5C). These probes consist of modified nucleotides with a methylene group bridging the 2’-O and 4’-C position of ribose ring, which converts the ribose ring into a locked conformation to enhance their stability and binding affinity, which makes it costlier. Based on the results, it’s evident that there is no necessity to opt for more expensive alternatives like LNA probes, as the DNA probes utilized in this method demonstrated comparable efficacy. This makes it a cost-effective approach for detecting small RNAs.

### Comparison of 3’end and 5’end biotin labeling of 5s rRNA

We also compared the sensitivity of 3’ end and 5’end biotin labelled DNA probes. A sequence specific probe against 5s rRNA was designed. In house probe labelling was done at the 3’end, while the 5’ end biotin labelled probe was commercially purchased. Detection was performed using a blot initially hybridized with the 3’ end-labeled probe, followed by stripping and subsequent hybridization with the 5’ end-labeled probe. Our results demonstrate more pronounced bands with the 3’ end-labeled probes in comparison to the 5’ end-labeled probe (figure 6), indicating better sensitivity of the 3’ end-labeled probes over the 5’ end-labeling.

**Figure 6:**
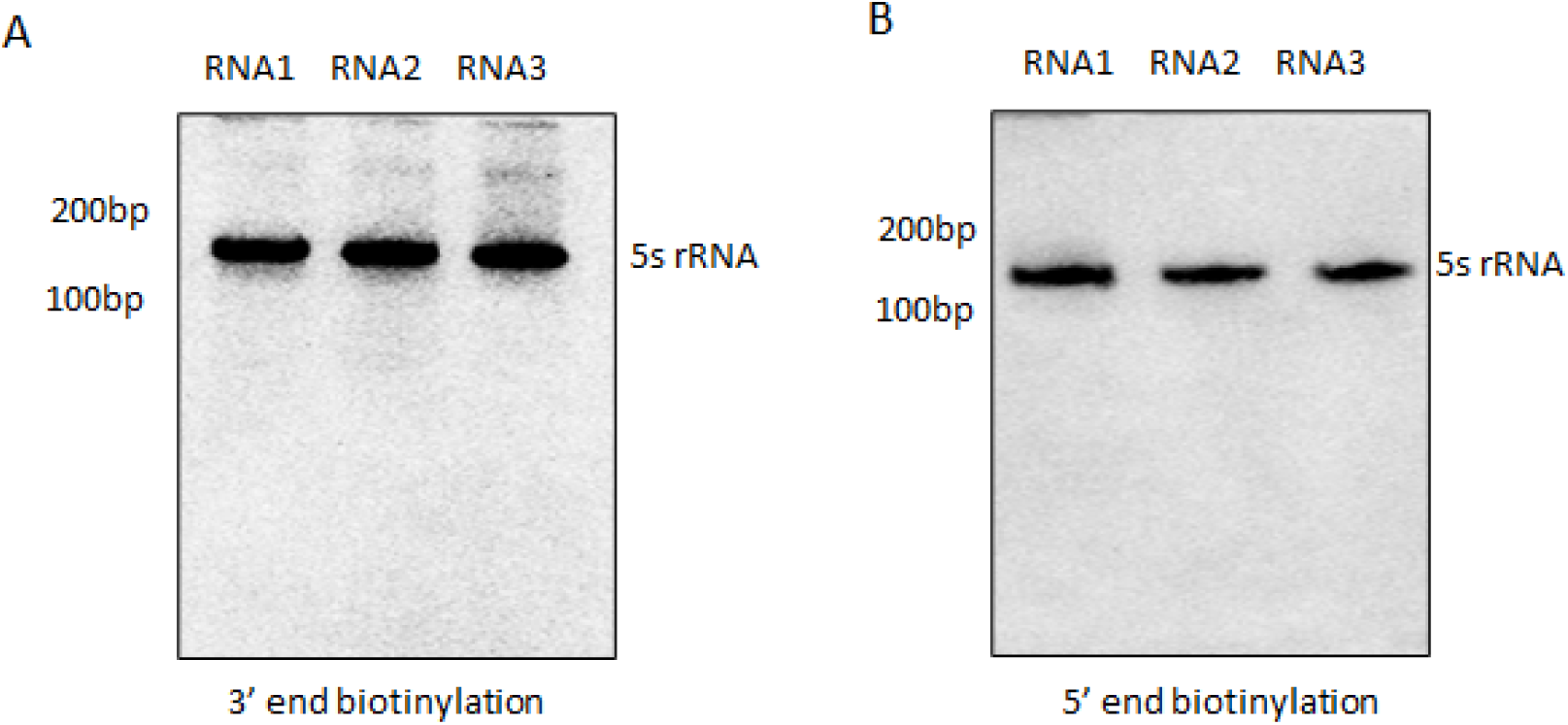
A comparative analysis between 3’ end and 5’ end biotinylation using the BiNoB technique: RNA samples (5 µg) were loaded onto 12% urea polyacrylamide gel electrophoresis (PAGE). The membrane was first hybridized with 3’ end biotin labelled 5s rRNA probe (left). Following stripping, the same membrane was rehybridized with 5’ end biotinylated probe (right).

### Small non-coding RNA (sncRNA) detection at different RNA concentration

The concentration of RNA often presents a limitation in the detection of small non-coding RNAs (sncRNAs). Previous studies have recommended using a minimum of 10-20 µg of RNA for sncRNA detection, making the Northern blotting technique challenging, particularly when working with samples containing limited RNA, such as blood samples. To address this limitation, we evaluated the minimum RNA amount required for efficient sncRNA detection. We tested various RNA quantities ranging from 250 ng to 2 µg. Using a probe targeting a 45-nt sncRNA tRNA fragment (ZZJQJYSWRYVMMV5BU6) derived from glutamine tRNA, we observed hybridization with both the full-length tRNA and its 45-nt fragment. At 250 ng, a faint tRNA band was visible, but no tRNA fragment band was detected. However, at 500 ng, the tRNA fragment began to appear with low intensity, while the tRNA band remained clearly visible. From 750 ng onwards, robust intensity bands for both the tRNA and its fragment were distinctly evident. These results support our finding that a total RNA quantity within the range of 500-750 ng is more than sufficient for comprehensive analysis of small non-coding RNAs (figure 7).

**Figure 7:**
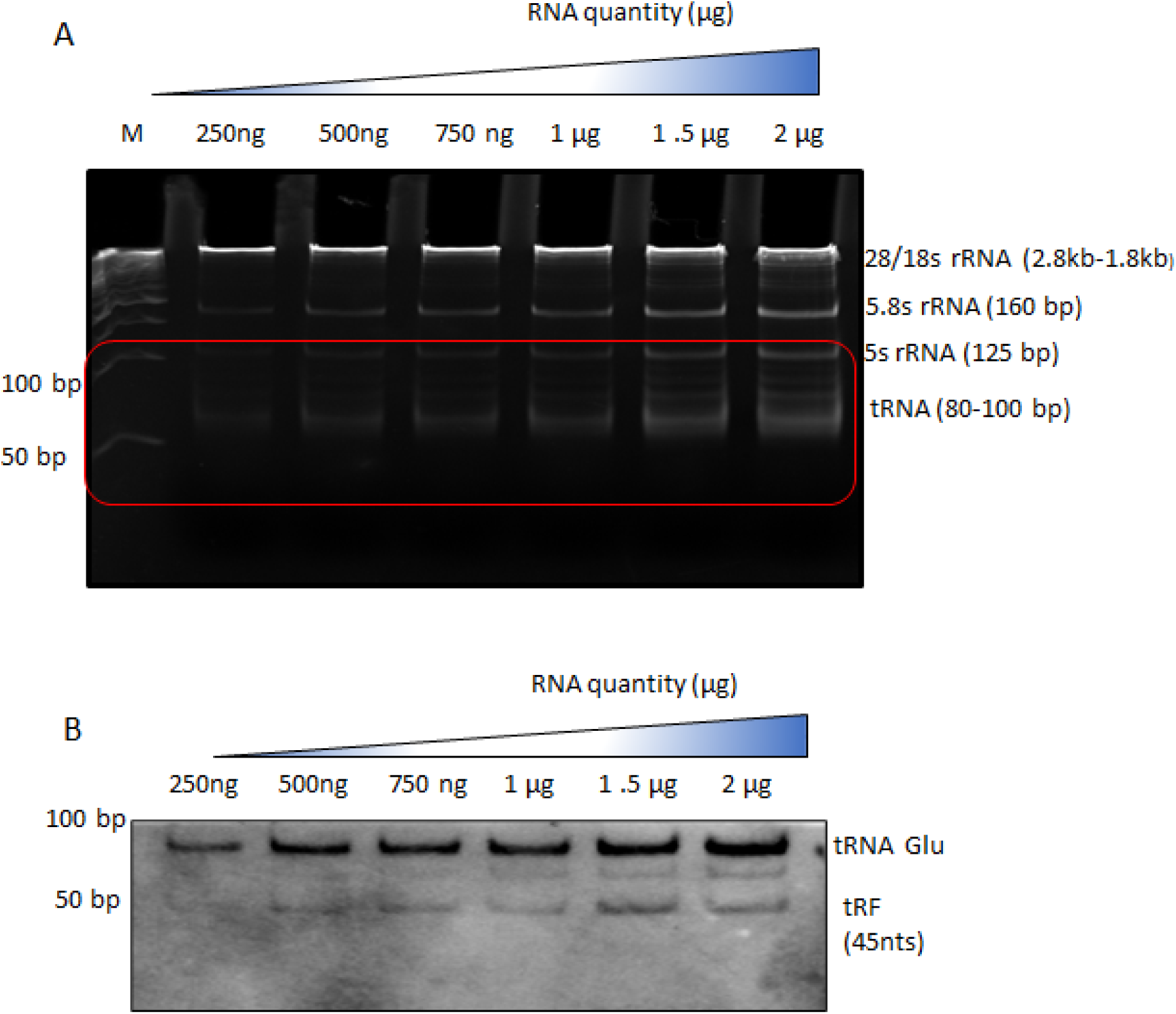
Detection of small non-coding RNAs at varying RNA quantities using BiNoB: Total RNA isolated from Huh7 cells was subjected to BiNoB technique with RNA quantities ranging from 250 ng to 2µg. The blot was hybridized with a 3’end biotinylated small non-coding-RNA i.e. tRNA fragment (9NZ6V6Z3M8ZLSSXUOLD2). The gel segment highlighted in red box (upper panel) has been shown in the blot (lower panel).

## Discussion

Northern blotting is considered as the most reliable method for analyzing the differential expression of the RNA transcripts. In this study, we have optimized the non-radioactive approach of northern Blotting termed as **<u>Bi</u>**otin based **<u>No</u>**rthern **<u>B</u>**lotting (BiNoB). The procedure overview has been depicted in figure 8 and detailed methodology are mentioned in supplementary 1. This approach is a one for all approach that can be used to detect large RNAs spanning kilo base pairs in lengths or small RNAs as short as 22 ntd long sncRNA. This technique eliminates the need for specialized set-ups, specialized training and is cost effective. Additionally, this approach also guarantees reproducible results with high resolution.

**Figure 8:**
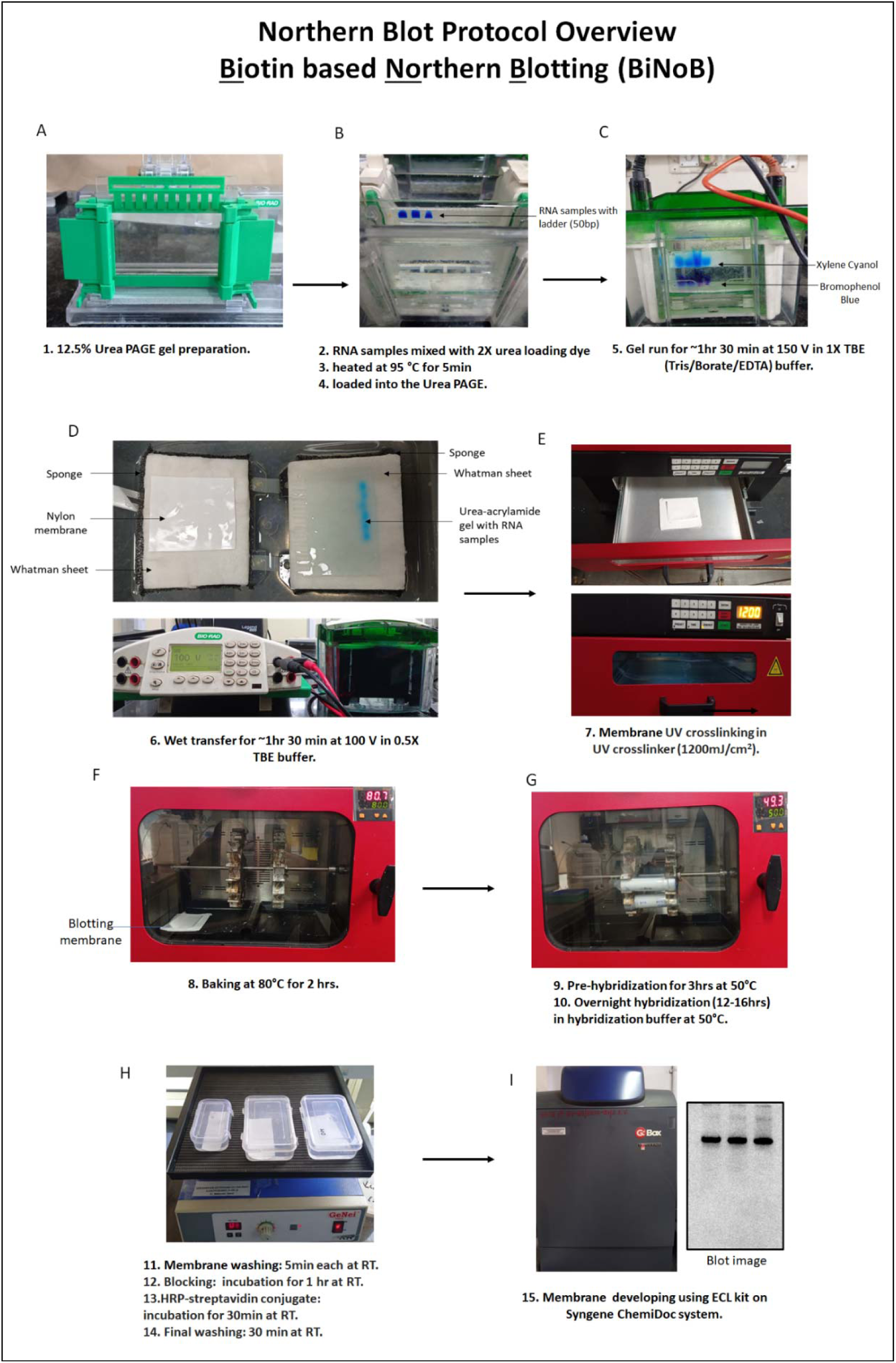
The BiNoB procedure overview.

The process involves vertical gel electrophoresis utilizing a wet transfer system, which ensures more effective transfer compared to the semi-dry method. This approach not only prevents gel overheating or breakage but also enhances the detection sensitivity of RNAs. The probes utilized are DNA oligos, offering a cost-effective advantage over LNA or RNA probes. These DNA probes are labeled with biotin 11-dUTP at their 3’ end through terminal deoxynucleotidyl transferase (TdT enzyme). This in-house probe labeling presents a cost-effective alternative to ordering pre-labeled synthetic probes. To enhance crosslinking efficiency, we have employed UV crosslinking at 1250KJ followed by baking at 80°C for 2 hours. Biotin labeling requires a secondary incubation with Streptavidin-HRP conjugate, facilitating easier detection using a Luminol based ECL kit and visualization on ChemiDoc system. Table 2 provides an overview of the distinctions between the traditional northern blotting method and the BiNoB technique.

We performed the membrane hybridizations after dot blot analysis to ensure efficient probe labelling. In this study, we detected varied lengths of RNAs, including 28s, 18s, and tubulin for long-sized RNAs exceeding 4 Kb (figure 3). Mid-sized RNAs, approximately 100 bp in length, such as 5S, U6 snRNA, and tRNAs, were also detected (figure 4). Small-sized RNAs, specifically miRNAs and tRNA fragments (tRFs) ranging from 20 to 50 nucleotides, were observed with enhanced resolution (figure 5). Notably, the method enabled the detection of parent tRNAs along with the associated tRFs generated as a result of tRNA fragmentation, which occurs through the action of various RNases. Furthermore, the approach facilitated the detection of three forms of miRNA: primary miRNA (1000 nts), precursor miRNA (150 nts), and miRNA ranging from 22 to 10 nts (figure5). This method holds promise for diverse miRNA-focused investigations, allowing for a comprehensive exploration of both tRFs and miRNA biogenesis.

We have also analyzed the efficacy of DNA probes and the LNA probes. Both worked equally well to detect ∼22nt RNA (figure 5). Additionally, we investigated the sensitivity of 3’ or 5’ labelled probes. In-house 3’end biotin labelling of DNA oligos was performed, while 5’end biotin labelled DNA probes were commercially ordered. 3’ end labelled probes were found to have higher sensitivity than the 5’end DNA probes (figure 6).

Northern blot analysis typically requires a minimum of 10 µg of total RNA for the detection of small non-coding RNAs (sncRNA). However, this requirement may pose limitations when working with RNA extracted from blood or tissues. In consideration of this constraint, we systematically optimized the minimal total RNA quantity necessary for effective sncRNA detection. Various RNA quantities, specifically 250 ng, 500 ng, 750 ng, 1 µg, 1.5 µg, and 2 µg, were investigated. For hybridization, a probe targeting a 48-nt sncRNA tRNA fragment was used. This probe exhibited hybridization with both the full-length tRNA and its 48-nt fragment. At 250 ng, a faint tRNA band was visible, while no tRNA fragment band was observed. At 500 ng, the band corresponding to tRNA fragment began to appear with low intensity, but the tRNA band remained clearly visible. Starting from 750 ng, robust intensity bands for both the tRNA and its fragment were distinctly evident. This substantiates our finding that a total RNA quantity within the range of 500-750 ng is more than adequate for comprehensive analysis (figure 7). This is the first study to report non-radioactive Northern blotting with a reduced quantity of RNA for the detection of small RNAs. Previously, due to limitations in RNA concentration, PCR-based techniques were the sole viable alternative. However, with the refined approach outlined here, we can confidently rely on the gold standard technique of northern blotting.

## Data Availability

All data used to support the findings of this study are included within the article.

## Acknowledgment

The authors acknowledge the members of the Verma laboratory for their critical comments.

## Author contributions

D.M., RR., and B.V.: conceptualization and methodology, formal analysis, investigation, writing–original draft; B.V.: supervision; B.V.: funding acquisition.

## References

1. Tong, L., Xue, H., Xiong, L., Xiao, J. & Zhou, Y. Improved RT-PCR Assay to Quantitate the Pri-, Pre-, and Mature microRNAs with Higher Efficiency and Accuracy. Mol Biotechnol (2015) doi:10.1007/s12033-015-9885-y.

2. Castoldi, M., Schmidt, S., Benes, V., Hentze, M. W. & Muckenthaler, M. U. miChip: An array-based method for microRNA expression profiling using locked nucleic acid capture probes. Nat Protoc (2008) doi:10.1038/nprot.2008.4.

3. Castoldi, M. et al. A sensitive array for microRNA expression profiling (miChip) based on locked nucleic acids (LNA). RNA (2006) doi:10.1261/rna.2332406.

4. Chen, C. et al. Real-time quantification of microRNAs by stem-loop RT-PCR. Nucleic Acids Res 33, 1–9 (2005).

5. Madhry, D. et al. Various transcriptomic approaches and their applications to study small noncoding RNAs in dengue and other viruses. Integrated Omics Approaches to Infectious Diseases 195–220 (2021) doi:10.1007/978-981-16-0691-5_12.

6. Löw, R. Nonradioactive Northern Blotting of RNA. in *Nucleic Acid Protocols Handbook*, The (2003). doi:10.1385/1-59259-038-1:239.

7. Carlile, M. & Werner, A. Extraction and nonradioactive detection of small RNA molecules. Methods in Molecular Biology (2014) doi:10.1007/978-1-4939-0931-5_8.

8. Meltzer, J. C. et al. Nonradioactive northern blotting with biotinylated and digoxigenin-labeled RNA probes. Electrophoresis (1998) doi:10.1002/elps.1150190825.

9. Várallyay, É., Burgyán, J. & Havelda, Z. MicroRNA detection by northern blotting using locked nucleic acid probes. Nat Protoc (2008) doi:10.1038/nprot.2007.528.

10. Válóczi, A. et al. Sensitive and specific detection of microRNAs by northern blot analysis using LNA-modified oligonucleotide probes. Nucleic Acids Res (2004) doi:10.1093/nar/gnh171.

11. Várallyay, É., Burgyán, J. & Havelda, Z. Detection of microRNAs by Northern blot analyses using LNA probes. Methods (2007) doi:10.1016/j.ymeth.2007.04.004.

12. Pall, G. S., Codony-Servat, C., Byrne, J., Ritchie, L. & Hamilton, A. Carbodiimide-mediated cross-linking of RNA to nylon membranes improves the detection of siRNA, miRNA and piRNA by northern blot. Nucleic Acids Res (2007) doi:10.1093/nar/gkm112.

13. Beckmann, B. M., Grünweller, A., Weber, M. H. W. & Hartmann, R. K. Northern blot detection of endogenous small RNAs (∼14 nt) in bacterial total RNA extracts. Nucleic Acids Res (2010) doi:10.1093/nar/gkq437.

14. Pall, G. S. & Hamilton, A. J. Improved northern blot method for enhanced detection of small RNA. Nat Protoc (2008) doi:10.1038/nprot.2008.67.

15. Huang, Q., Mao, Z., Li, S., Hu, J. & Zhu, Y. A non-radioactive method for small RNA detection by northern blotting. Rice 7, (2014).

16. Huang, Q., Mao, Z., Li, S., Hu, J. & Zhu, Y. A non-radioactive method for small RNA detection by northern blotting. Rice 7, (2014).

17. Beckmann, B. M., Grünweller, A., Weber, M. H. W. & Hartmann, R. K. Northern blot detection of endogenous small RNAs (∼14 nt) in bacterial total RNA extracts. Nucleic Acids Res 38, (2010).

18. Ahmad, W., Gull, B., Baby, J. & Mustafa, F. A comprehensive analysis of northern versus liquid hybridization assays for mrnas, small rnas, and mirnas using a non-radiolabeled approach. Curr Issues Mol Biol 43, 457–484 (2021).

19. Koscianska, E. et al. Northern blotting analysis of microRNAs, their precursors and RNA interference triggers. BMC Mol Biol 12, (2011).

