## Supplementary data for "Biotin based Northern Blotting (BiNoB): A Robust and Efficient Alternative"

### MATERIALS

#### REAGENTS

##### Chemical reagents

- **TRIzol Reagent:** Invitrogen (Carlsbad, California, United States) **CAUTION:** It contains phenol and guanidinium thiocyanate. Handle carefully.
- **Chloroform:** Qualigens (Mumbai, India)
- **Absolute Ethanol:** Merck (Darmstadt,Germany)
- **Urea:** (Amresco, Cgenomix)
- **Acrylamide and Bisacrylamide (38:2):** (Amresco, Cgenomix) **CAUTION:** Neurotoxic. Handle carefully.
- **TEMED:** Thermo Fisher (Massachusetts, United States)
- **Ammonium per Sulphate:** SRLchem
- **Tris:** Sigma Aldrich (Missouri, United States)
- **Borate:** Sigma Aldrich (Missouri, United States)
- **EDTA:** Sigma Aldrich (Missouri, United States) **CAUTION:** Carcinogenic and Mutagenic. Handle carefully.
- **Ficol400:** Sigma Aldrich (Missouri, United States)
- **Deionized Formamide:** Thermo Fisher (Massachusetts, United States)
- **Streptavidin-horseradish Peroxidase Conjugate:** Amersham, GE Healthcare
- **Sodium Phosphate dibasic:** Sigma Aldrich (Missouri, United States)
- **Sodium chloride:** Sigma Aldrich (Missouri, United States)
- **Polyvinyl pyrrolidone:** Sigma Aldrich (Missouri, United States)
- **Bovine Serum Albumin Type V:** HI Media
- **Ribolock RNases inhibitor:** ThermoFisher (Massachusetts, United States)
- **50bp DNA ladder:** ThermoFisher (Massachusetts, United States)
- **Xylene cyanol:** Sigma Aldrich (Missouri, United States)
- **Buffer AW1 and BufferAW2 (Catno.19072):** Qiagen (Hilden, Germany)

- **ULTRAhyb$^{TM}$ Ultrasensitive Hybridization Buffer:** Thermo Fisher (Massachusetts, United States) (Ambion:AM8670)
- **Terminal Deoxynucleotidyl transferase (TDT):** Thermo scientific cat no. EP0161
- **Biotin -d-11-UTP:** Thermo scientific cat no. R0081
- **Hybond N+ Nylon membrane (Cytiva)**
- **ECL Reagent:** Thermo:32106

#### EQUIPMENT

- Vertical gel system (Bio-Rad)
- Gel plates (Bio-Rad)

#### REAGENT SETUP

- **12% Urea-denaturating Polyacrylamide Gel:** The gel composition prepared with 6ml of 20% acrylamide: bis-acrylamide (19:1) solution, 4ml of 45% urea buffer prepared in 1x TBE (Tris Borate EDTA) buffer, 50µl of 10% APS, and 10µl TEMED.
- **20X TBE:** For 1 L, dissolve 216 g of 1M Tris base in 1 L of $ddH_2O$, 110 g of 1M Boric acid, 80 ml of 0.05M EDTA and bring up the volume to 1L with ddH2O. Dissolve the constituents in 1L of double distilled water. The buffer was sterilized by autoclaving and then stored at RT.
- **6X Urea Loading Dye:** The dye can be stored at -20°C for further use.
- **In-House Prepared Buffer:** In-house prepared buffer was prepared with 10mg/ml of BSA, 0.5M sodium phosphate buffer (pH 7.2),1mM EDTA and 7% SDS in the autoclaved double distilled water.
- **ETBR working solution:** Dissolve 1 g ethidium bromide in 100 ml distilled water. Store in dark bottle or wrap in aluminum foil.

#### PROCEDURE

##### RNA Extraction TIMING: 45 minutes

1. Total cellular RNA was isolated from different samples using the TRIzol method. Briefly, 300µl of chloroform was added to 500µl of TRIzol extracts followed by thorough mixing by tubes inversion and incubation at RT for 10min.

2.The tubes were centrifuged at 13,000rpm for 10min at 4°C to collect the aqueous supernatant.

**CRITICAL STEP:** Isolate aqueous phase carefully contains RNA only. As interphase contain proteins which compromises RNA quality.

To this supernatant, 600 µl of chilled absolute ethanol was added, thoroughly mixed, and then passed through RNA columns at 8000 RPM for 30 seconds. The column was subsequently washed with AW1 and AW2 buffers (Qiagen). Following 1min of empty spin, the RNA was finally eluted in nuclease-free water.

#### RNA quality and integrity check TIMING: 1 hour

3. RNA concentration was measured using Nanodrop 1000 and 260/280 and 260/230 ratios were checked for RNA quality.

**PAUSE POINT:** RNA can be stored at -80°C for more than 1 years in laboratory till further use.

4. Quality analysis of RNA by Urea-PAGE gel electrophoresis followed by EtBr staining. RNA samples about 1µg from Huh7, HEK cells and Vero cells were loaded into gel to assess quality. EtBr staining revealed clear bands of five significant RNA species, with 28 and 18S RNAs (ranging from 2.8Kb to 1.8Kb in size) observed exclusively in the wells, followed by 5.8s rRNA (160 nucleotides), 5s rRNA (125 nucleotides), and tRNA (80–100 nucleotides).

#### Sample Preparation and Gel electrophoresis TIMING: 2-2.5 hours

5. RNA samples were mixed with 2x Urea-loading dye (90mM Tris Base, 90mM Boric acid, 2mM EDTA, 12% Ficoll, 7M Urea, 0.03 %Xylene cyanol and Bromophenol blue dye) in equal volume followed by heat treatment at 95°C for 5 minutes.

6. A 12% Urea-PAGE gel was prepared for both high molecular weight (-more than 2kb) and low molecular weight (40bp-22bp) RNA.

**PAUSE POINT:** Allow gels to set for 15-20 minutes before use. Gels can be stored in plastic bag at overnight in 1X TBE at room temperature and 4°C.

7. Flush the wells of urea gel with running buffer before loading of RNA samples. The gel was pre-run for 45 minutes at 150V in 1x TBE buffer.

**CRITICAL STEP:** Flushing the wells of gel is an important step as urea got accumulated in the visible layer over bottom of each well which effect the proper resolution of RNA. Pre-run of the gels is important in 1X TBE buffer.

8.Samples were then loaded and run the gel at 150V till the bromophenol dye reached lower in the gel. The gel was then stained with EtBr (0.05 µg/ml) for 15 minutes followed by its visualization on ChemiDoc system (Syngene G: Box system).

## Membrane transfer and crosslinking   TIMING: 3-3.5 hours

9.Presoak the whatmann paper, sponges and the Hybond N+ Nylon membrane (Cytiva) in the transfer buffer (0.5x TBE). Wet transfer was carried out for 1 hour 30 minutes at 100V in 0.5x TBE buffer in mini trans-blot cell Bio-Rad apparatus.

10.Following transfer, membrane was UV crosslinked in UV crosslinker by Analytikjena at 1250mJ/cm$^2$. Baking was performed for 2 hours at 80°C after UV crosslinking.

**PAUSE POINT:** Blots can be stored at -80°C for further use.

## Pre-hybridization and Hybridization   TIMING: 4-16 hours

11. Membranes were pre-hybridized for 3hrs at 50°C in in-house prepared buffer (10mg/ml of BSA, 0.5M sodium phosphate buffer (pH 7.2), 1mM EDTA,7% SDS) following which overnight hybridization was set up for 12-16hrs by adding the probes to the same buffer at 50°C. commercially available ULTRAHyb$^{TM}$ buffer (Ambion:AM8670) was also used for comparing the results in both the buffers.

## Membrane blocking and streptavidin binding and washing   TIMING: 2 hours

12.Following hybridization process, the membrane was first washed with low stringency buffer (2X SSC+ 0.1% SDS) then by high stringency buffer (0.1% SSC+0.1% SDS) for five minutes each.

13.  The membrane was then subjected to blocking solution (0.1% skimmed milk in 1X TBS with 0.1% tween 20) for 1hr at room temperature.

14.Next, the membrane was incubated with HRP-Streptavidin conjugate at RT for 30 minutes followed by a final washing of 30 minutes in washing solution (1X TBS + 0.1% SDS).

## Probe labeling and dot blot analysis   TIMING: 1.5-2 hours

15.DNA probes were biotinylated at 3'terminal using Biotin 11-dUTP (Thermo: R0081) and terminal deoxynucleotidyl transferase (TdT) enzyme (Thermoscientific cat no. EP0161) in the presence of CoCl2 containing buffer.

16.The reaction was incubated at 37°C for 15 min followed by heat inactivation at 70°C for 10 min. After completion of the reaction, the biotinylated probe mixture was passed through a G-25 spin column (Cytiva) by centrifugation at 13,000 rpm for 1 minute at room temperature to remove unincorporated biotin moiety.

**PAUSE POINT:** Biotin labelled probes can be further stored at -20°C.

17. 5'end biotinylated probes were commercially purchased from Bioserve Biotechnologies (India) Pvt.Ltd to compare the sensitivity of both the probes. The sequences of probes used in the study has been mention in the table 2.

**PAUSE POINT:** 5'Biotin labelled probes is light sensitive and stored at -20°C for further use.

18.Biotinylation of probes was confirmed through dot blot analysis. Two microliter of each biotinylated probe was spotted onto the pre-soaked Hybond N+ Nylon membrane in 1x TBE buffer.

19.The membrane was then UV-cross-linked at (1250 mJ/cm2) followed by incubation in blocking solution (0.1% skimmed milk in 1X TBS with 0.1% tween 20) for 30 minutes at RT.

20.The membrane is finally incubated with streptavidin-HRP conjugate at RT for 20 minutes followed by washing with low-stringency buffer (2X SSC + 0.1% SDS) and high-stringency buffer (0.1% SSC + 0.1% SDS) for 5 minutes each.

#### Signal detection TIMING: 10 minutes

21.After final washing step, the blot was finally developed using ECL reagents (Thermo:32106) and the blot was finally visualized on ChemiDoc system.
